# Bcl-xL is translocated to the nucleus via CtBP2 to epigenetically promote metastasis

**DOI:** 10.1101/2023.04.26.538373

**Authors:** Tiantian Zhang, Sha Li, Yingcai Adrian Tan, Joseph HyungJoon Na, Zhengming Chen, Priyadarshan Damle, Xiang Chen, Soyoung Choi, Bikash Mishra, Dunrui Wang, Steven R. Grossman, Xuejun Jiang, Yi Li, Yao-Tseng Chen, Jenny Z. Xiang, Yi-Chieh Nancy Du

## Abstract

Besides its mitochondria-based anti-apoptotic role, Bcl-xL also travels to the nucleus to promote cancer metastasis by upregulating global histone H3 trimethyl Lys4 (H3K4me3) and TGFβ transcription. How Bcl-xL is translocated into the nucleus and how nuclear Bcl-xL regulates H3K4me3 modification are not understood. Here, we report that C-terminal Binding Protein 2 (CtBP2) binds Bcl-xL via its N-terminus and translocates Bcl-xL into the nucleus. Knockdown of CtBP2 by shRNA decreases the nuclear portion of Bcl-xL and reverses Bcl-xL-induced cell migration and metastasis in mouse models. Furthermore, knockout of CtBP2 suppresses Bcl-xL transcription. The binding between Bcl-xL and CtBP2 is required for their interaction with MLL1, a histone H3K4 methyltransferase. Pharmacologic inhibition of MLL1 enzymatic activity reverses Bcl-xL-induced H3K4me3 and TGFβ mRNA upregulation as well as cell invasion. Moreover, cleavage under targets and release using nuclease (CUT&RUN) coupled with next generation sequencing reveals that H3K4me3 modifications are particularly enriched in the promotor region of genes encoding TGFβ and its signaling pathway in the cancer cells overexpressing Bcl-xL. Altogether, the metastatic function of Bcl-xL is mediated by its interaction with CtBP2 and MLL1.

## Introduction

Bcl-xL has long been known for its anti-apoptotic function during embryonic development and in pathological conditions (Boise *et al*, 1993). Bcl-xL executes its anti-apoptotic function in the mitochondrial membrane by binding to and inhibiting the pro-apoptotic activity of Bax/Bak, which are otherwise poised to initiate the apoptotic cell death pathway. Bcl-xL is frequently overexpressed in cancer, whether newly diagnosed or resistant to therapies (Hafezi & Rahmani, 2021). Moreover, the role for Bcl-xL in tumor metastasis has been previously ascribed to its anti-apoptotic function, e.g., Bcl-xL increases metastasis by providing a survival advantage to the tumor cells (Fernandez *et al*, 2000).

We have previously investigated the interdependence of Bcl-xL’s metastatic function and its canonical anti-apoptotic function. Using multiple mutants, cell lines, and mouse models, we have discovered that Bcl-xL promotes metastasis independent of its canonical anti-apoptotic function and that the metastatic function requires its nuclear translocation (Choi *et al*, 2016; Du *et al*, 2007). We have demonstrated that ABT-737, a prototype Bcl-xL inhibitor, does not affect the migration function of Bcl-xL (Choi *et al*., 2016). Similar to wild-type (wt) Bcl-xL, anti-apoptosis-defective Bcl-xL mutants and an engineered Bcl-xL targeted to the nucleus can still promote tumor cell migration and invasion (Choi *et al*., 2016). In contrast, engineered Bcl-xL – either targeted to the mitochondria or excluded from the nucleus – does not promote cell migration and invasion even though they protect cells from apoptosis (Choi *et al*., 2016). Importantly, nuclear Bcl-xL are detected in patients’ liver metastases of pancreatic neuroendocrine tumors (PNET) and in the mouse models of human PNET xenografts (Choi *et al*., 2016). Additionally, a patient-associated Bcl-xL mutation, N136K, disrupts its anti-apoptotic function but promotes breast cancer cell migration (Zhang *et al*, 2020). Overexpressed Bcl-xL increases secreted Transforming growth factor β (TGFβ) (Choi *et al*., 2016; Weiler *et al*, 2006), which is widely known to promote cancer metastasis (Massague, 2008).

As the C-terminal transmembrane domain of Bcl-xL prevents its nuclear import (Kaufmann *et al*, 2003), overexpressed Bcl-xL proteins in cancer cells cannot by themselves freely enter the nucleus. Therefore, the mechanism of Bcl-xL’s nuclear translocation is unknown. In this study, we identify Bcl-xL-interacting proteins that promote its nuclear translocation for executing its metastatic function.

## Materials and Methods

### Immunofluorescent analysis

Sections of formalin-fixed paraffin-embedded primary breast cancer tissues and lymph node metastases were deparaffinized and rehydrated by passage through a graded xylene/ethanol series before staining. Cells were cultured on poly-D-lysine (PDL)-coated glass coverslips for 24 hours before fixation in 4% paraformaldehyde in PBS for 10 min. After permeabilizing and blocking in 0.5% BSA with PBS plus 0.025 % Triton X-100, 0.02% NaN_3_ and 0.3 mM DAPI for 1 hours, coverslips were incubated with primary antibodies (mouse anti-HA 1:200, Cell Signaling Technology, 2367; mouse anti-CtBP2, 1:250; BD Biosciences Cat# 612044, RRID:AB_399431); rabbit anti-Bcl-xL 1:200, Cell Signaling Technology, 2764s; MTCO1, 1:500; Abcam, ab14705) overnight at 4°C. The coverslips were then washed three times with PBS for 15 min, followed by incubation with Alexa Fluor® 647 goat anti-mouse (1:500; Life Technologies, A21236), Alexa Fluor® 488 goat anti-mouse (1:400, Life Technologies), or Alexa Fluor® 647 goat anti-rabbit (1:400, Life Technologies) at room temperature in the dark for 2 hours, and were washed three times with PBS for 15 min. Coverslips were mounted with Vectashield mounting medium (Vector Labs) and were examined using confocal microscopy (Olympus FLUOVIEW FV10i). The total, nuclear, and cytoplasmic immunofluorescence signals of more than 30 cells per cell line were quantified using ImageJ (ImageJ, RRID:SCR_003070).

### Sequencing of genomic BCL2L1 (Bcl-xL)

Genomic DNA was extracted from breast cancer patient lymph node metastases using the DNeasy Blood & Tissue Kit (QIAGEN, 69504). Exon 1, 2, and 3 of BCL2L1 were amplified by PCR and cloned into pCR-Blunt-TOPO vector using Zero Blunt TOPO PCR Cloning Kit (Thermo Fisher Scientific Cat# 450245) The primers used for PCR are exon 1&2 F1: AACTCTTCCGGGATGGGGTAA; exon 1&2 R1: CCAGCCGCCGTTCTCCT; exon 1&2 F2: ATGGCAGCAGTAAAGCAAGCG; exon 1&2 R2: CCCATCCCGGAAGAGTTCATTCA; exon 1&2 F3: AGAAGGGACTGAATCGGAGATG; exon 1&2 R3: CGAAGGAGAAAAAGGCCACAAT; exon 3 F1: GATACTTTTGTGGAACTCTA; and exon 3 R1: GGTAGAGTGGATGGTCAGT. M13 forward primer GTAAAACGACGGCCAG was used for sequencing.

### Reversible cross-link immuno-precipitation (ReCLIP)

10-cm dishes of 90% confluent cells were used and cells were washed three times with cold PBS to remove all traces of media. Following removal of PBS, 5 mL of a 1 mM DSP solution in PBS was added to each dish. Cells were incubated for 30 min at 37°C, 5% CO_2_ with occasional swirl. Crosslinker solution was then removed, and 5 mL of 50 mM Tris (pH 8.0) was added to each plate for an additional 10 min to quench the crosslinker. Following quenching, plates were placed on ice bath and washed one more time with cold PBS and lysed with freshly-prepared CLIP buffer (50 mM Tris (pH 8.0), 150 mM NaCl, 1 mM EDTA, 0.1% SDS, 0.5% Sodium Doxycholate, 20 mM HEPES, 1% Triton X-100) plus protease inhibitors and phosphatase inhibitors. Lysates were homogenized by end-over-end rotation for 30 min at 4°C and cleared by centrifugation. Clarified lysates were incubated with anti-HA Magnetic Beads (Thermo Scientific, 88836) or anti-HA agarose (Thermo Scientific, 26181) for 2 hours at 4°C with end-over-end rotation. The beads were then washed 5 times with 1 mL CLIP buffer supplemented with protease and phosphatase inhibitors. Proteins were eluted by incubating the beads with 2X loading buffer (without DTT) at 50°C with gently shaking for 10 min. Eluted proteins were isolated by centrifugation, DTT was added to a final concentration of 100 mM and samples were boiled for 3 min.

### Western blot

Whole cell lysates were prepared by lysing cells with RIPA buffer (0.1% SDS, 1% Triton X-100, 0.5% sodium deoxycholate, 25 mM Tris (pH 8.0), 150 mM NaCl, 1 mM EDTA) supplemented with a protease inhibitor mixture and PhosSTOP (Roche). Histones were prepared by acid extraction as previously described (Choi *et al*., 2016). Proteins were quantified with Bradford assay (Bio-Rad). Equal amounts of proteins were separated with SDS-PAGE and transferred to nitrocellulose membranes. To visualize equal protein loading, blots were stained with Ponceau S. Blots were incubated in 5% non-fat milk in TBST, probed with primary antibodies to HA (1:1,000, Cell Signaling Technology, Cat# 2367), CtBP2 (1:1000, BD Biosciences, Cat# 612044, RRID:AB_399431), Mek1/2 (1:1,000, Cell Signaling Technology, Cat# 9122), alpha-tubulin (Sigma, Cat# T5168), MLL1 (1:1,000, Cell Signaling Technology, Cat# 14197), CtBP1 (1:1,000, BD Biosciences, Cat# 612042, RRID:AB_399431), Myc tag (1:1,000, Cell Signaling Technology, Cat# 2276), V5 (1:1,000, Invitrogen, Cat# R950-25), Histone H3K4me3 (1:1,000, Cell Signaling Technology, Cat# 9751, RRID:AB_2616028), and Histone 3 (1:1,000, Cell Signaling Technology, Cat# 5427) and then were incubated with horseradish peroxidase-conjugated secondary antibodies. Protein bands were visualized by enhanced chemical luminescence (Cytiva or Pierce).

### Generation of cell lines expressing inducible shRNAs

The shRNAs for human CtBP2 and MLL1 were designed using the SplashRNA algorithm (Pelossof *et al*, 2017) and cloned into the LT3RENIR vector, expressing dsRed fluorescence-coupled miR-E shRNAs from an optimized Tet-responsive element promoter (T3G) (Fellmann *et al*, 2013). Four dox-inducible shRNA against different regions of human CtBP2 (tet-O-dsRed-shCtBP2-PGK-rtTA3 (TR-shCtBP2) and one control shRNA against Renila Luciferase (shRLuc #713) were constructed within the lentiviral miRE vector (Fellmann *et al*., 2013) by MSKCC Gene Editing & Screening Core. Antisense guide sequences of these shRNAs are listed in Supplementary Table S2. DNAs were transfected into 293T cells using *Trans*IT®-Lenti Transfection Reagent (Mirus, MIR2304). Viral supernatants were collected, filtered, and used to infect BON1/TGL cells in the presence of 8 µg/ml polybrene. Cells were selected with 0.8 mg/ml G418 (Corning, 108321-42-2) 72 hours after infection for 12 days, and then stable cell lines were maintained in 0.4 mg/ml G418. For fluorescence-activated cell sorting (FACS), BON1/HA-Bcl-xL/TR-shRLuc #713, BON1/HA-Bcl-xL/TR-shCtBP2 #2260, and BON1/HA-Bcl-xL/TR-shCtBP2 #2403 cells were pre-treated with 0.5 μg/ml doxycycline for 96 hours and sorted using a BD FACSAria II Cell Sorter System at Weill Cornell Medicine Flow Cytometry Core Facility. All the parental cell lines were authenticated by University of Arizona Genetics Core before use.

### DNA, cell lines, and transient transfection

U2OS, 293T, N134, MDA-MB-231 (RRID:CVCL_ZZ22), and MDA-MB-231/CtBP2 knock out (KO) cells were maintained in Dulbecco’s modified Eagle’s medium (DMEM) supplemented with 10% FBS, 0.2 mM L-glutamine, 100 units/ml penicillin and 100 µg/ml streptomycin. BON1/TGL/pQCXIP (pQ, vector control) and BON1/TGL/HA-Bcl-xL (wt or mutants) cells have been described (Choi *et al*., 2016) and maintained in above medium with 0.5 μg/ml puromycin. Cells were cultured at 37°C in a standard tissue culture humidified chamber.

V5-tagged CtBP2 constructs (Full length (445 aa), residues 1 to 321, and residues 322 to 445) were described in (Paliwal *et al*, 2006) and Bcl-xL/Bcl-B chimeric constructs was described in (Saurabh *et al*, 2014). The plasmids for the deletion mutants (CtBP2-V5-mut1: 33-321 aa, CtBP2-V5-mut2: 61-321 aa, CtBP2-V5-mut3: 82-321 aa, and CtBP2-V5-mut4: 105-321 aa) were constructed using N-terminal CtBP2 (1-321aa) as template. The mutants were PCR amplified by round-the-horn amplification method. The oligos used are listed in Supplementary Table S3. The linear DNA was resolved on a 1% agarose gel, extracted, and circularized by Gibson assembly. All the constructs were confirmed by sequencing. The MDA-MB-231 cell line with CtBP2 knockout was generated using a previously cloned CRISPR-Cas9 plasmid (Chawla *et al*, 2018). Briefly, MDA-MB-231 cells at 70% confluency were transfected with the CRISPR-Cas9-CtBP2 plasmid. After 72 hours, the cells were single sorted on an Aria-BD, (San Jose, CA, USA) FACSAria™ II High-Speed Cell Sorter (λ exc =488nm) in 96-well plates. The cells were grown up to confluency and then replica plated in other 96-well plates. These clones were screened by Western blotting to identify homozygous knockout of CtBP2.

The following amount of DNA was used in transient transfection into U2OS cells: 1 μg HA-Bcl-xL, 1 μg pQ vector, 1μg full length-CtBP2-V5, 2 μg N-terminal CtBP2-V5, 8 μg C-terminal CtBP2-V5, 2 μg CtBP2-V5-mut1, 2 μg CtBP2-V5-mut2, 4 μg CtBP2-V5-mut3, and 6 μg CtBP2-V5-mut4. The following amount of DNA was used in transient transfection into 293T cells: 2 μg full length-CtBP2-V5, 2 μg either Myc-Bcl-xL, Myc-Bcl-B, Myc-Bcl-B/xL(BH1-BH2-TM), Myc-Bcl-xL/B(BH4-loop), Myc-Bcl-xL/B(loop), Myc-Bcl-B/xL(loop), Myc-Bcl-xL/B(BH1-BH2-TM), Myc-Bcl-B/xL(BH4-loop). The plasmids were incubated in 200 μl Opti-MEN medium mixed with X-tremeGENE HP DNA Transfection Reagent (Sigma) (ratio of reagent: DNA = 3 μl : 1 μg) at room temperature for 30 min. Cells at a confluence of 50∼60% were used for each transfection. Cells were washed with PBS, then DMEM containing 1% FBS and 0.2 mM L-glutamine (without antibiotics) were added. Transfection complexes were added to cells in a dropwise manner and the dishes were gently swirled. Cells were incubated for 48 hours at 37°C, 5% CO_2_ before protein lysates were made. All the parental cell lines were authenticated by University of Arizona Genetics Core before use.

### Generation of N134 cell lines expressing Bcl-xL/Bcl-B chimeric constructs

Bcl-xL/B(BH4-loop) and BclB/xL(BH4-loop) were PCR amplified from Bcl-xL/Bcl-B chimeric constructs (Saurabh *et al*., 2014), subcloned into pENTR3C-HA-Bcl-xL, and then recombined into RCASBP-Y-DEST (Loftus *et al*, 2001) using LR Clonase (Invitrogen) according to the manufacture’s instruction. Constructs were confirmed by DNA sequencing. Generation of N134 cell lines expressing these RCASBP constructs was based on the method previously described (Du *et al*., 2007; Zhang *et al*, 2017).

### Reverse transcription quantitative real-time PCR (RT-qPCR)

Cell lines (MDA-MB-231, MDA-MB-231 with CtBP2 KO, BON1/TGL/pQ and BON1/TGL/HA-Bcl-xL) grown on 6-cm dishes were used to isolate mRNA using RNeasy Plus Mini kit (Qiagen, 74136) containing gDNA Eliminator spin columns. cDNA was generated using the SuperScript™ IV VILO™ Master Mix with ezDNase™ Enzyme (Invitrogen, 11766050), and power SYBR green-based quantitative real-time PCR was performed using primers specific for human *BCL2L1* (forward: 5′- GGTCGCATTGTGGCCTTT -3′, reverse: 5′- TCCGACTCACCAATACCTGCAT - 3′), human *CtBP2* (forward: 5′- GAGAGTGATCGTGCGGATAGG -3′, reverse: 5′- TTGCACACGGCAATTCCGA -3′), human *ACVR1* (forward: 5′- GCGGTAATGAGGACCACTGT -3′, reverse: 5′- CCCTGCTCATAAACCTGGAA -3′), human *SMAD3* (forward: 5′- GCCTGTGCTGGAACATCATC-3′, reverse: 5′- TTGCCCTCATGTGTGCTCTT -3′), human *NKX2-5*(forward: 5′- CCAAGTGCTCTCCTGCTTTCC -3′, reverse: 5′- CGCGCACAGCTCTTTTTTATC - 3′), human *SMAD5* (forward: 5′- CTGGGATTACAGGACTTGACC -3′, reverse: 5′- AAGTTCCAATTAAAAAGGGAGGA -3′), human *ZNF423* (forward: 5′- GGAACAGCGTGACAAGTCAAG -3′, reverse: 5′- ACAGTGATCGCAGGTGTAAATTG - 3′), human *MRPL19* (forward: 5′-GGGATTTGCATTCAGAGATCAGG -3′, reverse: 5′- CTCCTGGACCCGAGGATTATAA -3′), human *HMBS* (forward: 5′- CCATCATCCTGGCAACAGCT -3′, reverse: 5′-GCATTCCTCAGGGTGCAGG -3′), with the comparative CT method (ΔΔCT; Applied Biosystems™ 7500 Real-Time PCR). Both *MRPL19* and *HMBS* were used together as endogenous controls for normalization.

### Animal experiments

NSG mice were purchased from the Jackson Laboratory. All mice were housed in accordance with the institutional guidelines. All procedures involving mice were approved by the institutional animal care and use committee of Weill Cornell Medicine. NSG mice at the age of ∼10 weeks were fed with dox-containing diet (Envigo, TD.00426) one day before the injection of cells and injected with one million cells that were treated with doxycycline (0.5 μg/ml for 96 hours) in 100 μl PBS via the left ventricle. For tail vein metastasis assay, 1.5 × 10^6^ N134 cells in 150 μl PBS were injected into the tail veins of NSG mice at the age of ∼10 weeks. Mice were subjected to bioluminescent imaging using the *In Vivo* Imaging System Spectrum (PerkinElmer) as described (Choi *et al*., 2016; Choi *et al*, 2019; Du *et al*, 2011; Du *et al*., 2009).

### *In vitro* transwell invasion assay

BON1/HA-Bcl-xL/TR-shCtBP2 and BON1/HA-Bcl-xL/TR-shRLuc #713 cells were treated with 1 µg/ml (before sorting) or 0.5 µg/ml (after sorting) doxycycline (Clontech Laboratories, Inc., 631311) for 96 hours, and 2 × 107 of cells were seeded in Matrigel-coated transwell invasion chambers (Corning, 354480) with DMEM containing 1% FBS, 0.4 mg/ml G418, 0.5 µg/ml puromycin, 0.2 mM L-glutamine and 100 units/ml penicillin and 100 µg/ml streptomycin. The lower chambers were filled with DMEM containing 10% FBS, 0.4 mg/ml G418, 0.2 mM L-glutamine and 100 units/ml penicillin and 100 µg/ml streptomycin. Fifty thousand cells per cell line were seeded in 1% FBS, 0.4 mg/ml G418, 0.5 µg/ml puromycin, 0.2 mM L-glutamine and 100 units/ml penicillin and 100 µg/ml streptomycin per well of 24-well dish for the proliferation controls. After 24 hours of incubation, cells on the opposite side of the chambers were fixed in 4% paraformaldehyde in PBS for 10 min, washed with PBS-/- for 5 min, stained with 0.1% crystal violet for 30 min and counted in 8 fields under 20X magnification.

### Subcellular fractionation

Subcellular fractionation was performed based on the method previously described (Choi *et al*., 2016).

### Caspase-3/-7 activity assay

A total of 1 × 10^4^ cells (N134 parental cells and N134 expressing wt Bcl-xL, HA-Bcl-B/xL(BH4-loop) (construct #5), or RCASBP–HA-Bcl-xL/B(BH4-loop) (construct #6)) were seeded in 100 µL culture medium in a 96-well white wall plate. After overnight incubation, cells were treated with DMSO or 10 mM etoposide for 24 hours. Caspase-3 and -7 activities were measured using the Caspase-Glo 3/7 assay kit (Promega, G8090) according to manufacturer’s instructions. Luminescence was measured using a Glomax Multi Plus Detection System (Promega, Madison, WI, USA), and caspase 3/7 activities was normalized by the number of cells. Data represented 3 replicates per condition and were expressed as mean of fold change over N134 cells without etoposide ± SEM.

### 3D tumor sphere invasion assay

100 BON1/HA-Bcl-xL/TR-shMLL1 #13408, BON1/HA-Bcl-xL/TR-shMLL1 #13406, and control BON1/HA-Bcl-xL/TR-shRLuc #713 cells in 100 µl growth medium were seeded into 96-well round bottom ultra-low attachment plates (Corning, 4515) with 2.5% (v/v) Matrigel (Corning, 354234). Plates were incubated for 96 hours at 37 °C, 5% CO_2_, 95% humidity for formation of a single spheroid of cells. After 96 hours, spheroids were then treated with 0.5 μg/ml doxycycline in fresh medium containing Matrigel (Day 0). Fresh medium containing 0.5 μg/ml doxycycline was added every 48 hours. Images were taken at the indicated time points. 6 wells of each cell/condition were used and analyzed. Original images were under × 20 magnification.

### CUT&RUN and next generation sequencing analysis

BON1/TGL/pQ and BON1/TGL/HA-Bcl-xL cells were washed with cold PBS and scraped from cell culture dishes (triplicates). Cells were pelleted at 4°C, 3,000g for 5 minutes. CUT&RUN was performed from 500,000 cells per condition in triplicates as previously described in (Skene *et al*, 2018) using the following antibodies: rabbit anti-HA (Cell Signaling Technology, Cat# 3724); rabbit anti-MLL1 (Epicypher, Cat# 13-2004); rabbit anti-H3K4me3 (Cell Signaling Technology, Cat# 9751, RRID:AB_2616028); rabbit anti-mouse IgG (Abcam, ab46540). Briefly, cells were harvested and bound to concanavalin A-coated magnetic beads (Bangs Laboratories, BP531) after a 10 min incubation at room temperature on a rotator. Cell membranes were permeabilized with 0.0125% digitonin (Millipore-Sigma, #300410) and the different antibodies were incubated for 20 min at RT on a rotator. Beads were washed and incubated with pAG-MNase. The pAG-MNase plasmid was a gift from Steven Henikoff (RRID:Addgene_123461) and the pAG-MNase recombinant protein was expressed and purified as previously described in (Meers *et al*, 2019). CaCl_2_-induced digestion occurred on ice for 30 min and was stopped by chelation at 37°C for 20 min. Spike-in drosophila DNA (Zyagen, GD-290) was added in the chelation buffer at 50 pg per condition. DNA was finally isolated using the MinElute PCR Purification Kit (Qiagen, 28006) following the manufacturer’s instructions. After quantification of the recovered DNA fragments, libraries were prepared using the ThruPLEX®DNA-Seq kit (Takara, R400676) following the manufacturer’s instructions, purified with AMPure XP magnetic beads (Beckman Coulter, A63881), and quantified using a Qubit fluorometer (Thermo Fisher Scientific).

The quantity and quality of the pooled Cut & Run libraries were measured by Invitrogen Qubit Fluorometer (Thermo Fisher Scientific) and Agilent Tape Station with High Sensitivity D1000 Screen Tape. The average libraries size was 300 bp. According to the manufacturer’s instructions, denatured and diluted libraries were loaded to the bottom of the cartridge reservoir provided with the Illumina NextSeq 2000 reagents kit, and sequenced at paired-end 50 bps on NextSeq 2000 sequencer (Illumina). The raw sequencing reads in BCL format were processed through bcl2fastq 2.20 (Illumina) for FASTQ conversion and demultiplexing. Data were processed using Cut&RunTools (https://bitbucket.org/qzhudfci/cutruntools/src/default/) (Zhu *et al*, 2019) with the default settings. Briefly, reads were adapter trimmed using Trimmomatic (Trimmomatic, RRID: SCR_011848) (Bolger *et al*, 2014), and an additional trimming step was performed to remove up to 6 bp adapter from each read. Next, reads were aligned to the hg19 genome using bowtie 2 (Langmead & Salzberg, 2012) with the ‘dovetail’ settings enabled. Alignments were further divided into ≤ 120-bp and > 120-bp fractions. Alignments from the > 120-bp fractions were used for peak calling with MACS2 (2.1.1) (Zhang *et al*, 2008), followed by de novo motif searching within the peak regions with MEME suite (4.12.0) (Machanick & Bailey, 2011). Differential binding analysis was conducted with Bioconductor, DiffBind (Bioconductor, RRID:SCR_006442) (Ross-Innes *et al*, 2012). The list of 1,910 unique genes/promoters with H3K4me3 binding sites in HA-Bcl-xL overexpressing cells were input into Ingenuity Pathway Analysis (Ingenuity Pathway Analysis, RRID:SCR_008653) for pathway and network enrichment analysis. The peaks of each condition were merged using SAMTOOLS (SAMTOOLS, RRID:SCR_002105), indexed, and normalized into bigwig files for IGV browser.

### Statistical analysis

Radiance values of luciferase signals or 3D tumor spheroid sizes (area) were transformed to the natural log scale before analysis. Generalized Estimating Equations (GEE) method was used to test the overall difference in radiance signals or whole area change over time and the analysis were performed in statistical software Statistical Analysis System (Statistical Analysis System, RRID:SCR_008567). *In vitro* experiments were performed at least in triplicates.

## Results

### Nuclear Bcl-xL is detected in breast cancer patients

In healthy cells, most of the Bcl-xL proteins reside in the mitochondria, while some are localized in the cytosolic and ER (Popgeorgiev *et al*, 2018). However, we have found nuclear Bcl-xL in the liver metastases of pancreatic neuroendocrine tumor (PNET) by immunofluorescent staining, and our studies suggested that nuclear Bcl-xL promotes metastasis (Choi *et al*., 2016). To broaden the clinical relevance of nuclear Bcl-xL beyond PNET, we sought to determine whether nuclear Bcl-xL can also be found in specimens of breast cancer, the most common malignancy in women. We conducted immunofluorescent analysis using antibodies against Bcl-xL and MTCO1 (a mitochondrial marker) and visualized nuclear DNA with 4′,6-diamidino-2-phenylindole (DAPI). We detected predominant nuclear Bcl-xL in 3 of 15 cases of breast cancer specimens (Figure 1a, 1b, and data not shown) and weak nuclear Bcl-xL in the other cases (Figure 1c and data not shown), suggesting that Bcl-xL can also be translocated into the nucleus of the breast cancer cells.

**Figure 1.**
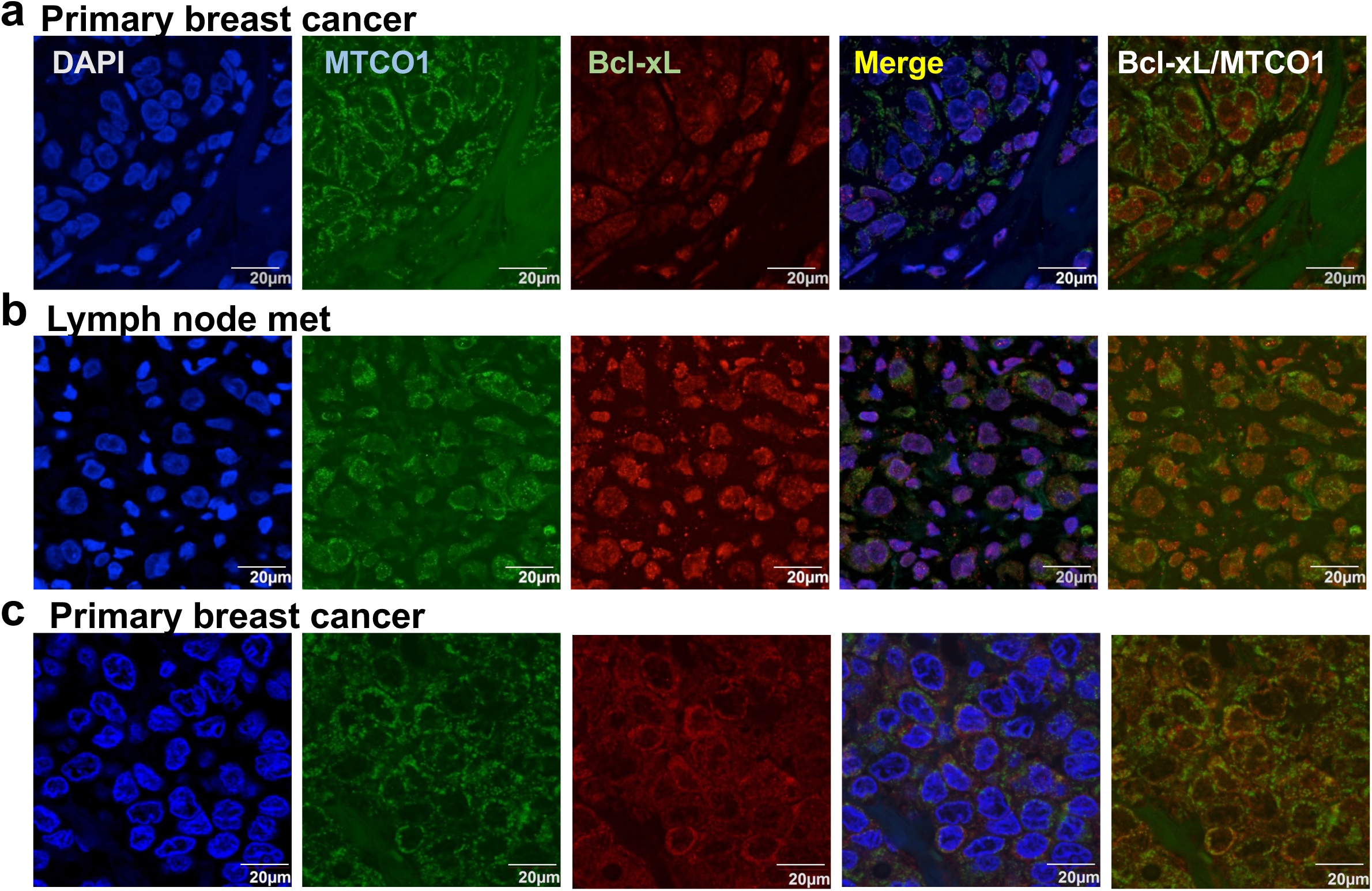
Nuclear Bcl-xL is found in human breast cancer specimen. Confocal microscopy images of Bcl-xL (red), mitochondrial marker MTCO1 (green), and DAPI (blue) from two primary human breast cancer (**a** and **c**) and lymph node met (**b**). Scale bar, 20 μm. Original magnification, × 60.

To determine whether mutations of Bcl-xL contribute to the nuclear translocation, we micro-dissected the breast cancer case that had mostly nuclear Bcl-xL, purified genomic DNA, PCR amplified Bcl-xL, and sequenced the entire Bcl-xL gene. We found no mutation in Bcl-xL (Supplementary Figure S1), suggesting that the change of subcellular localization of Bcl-xL in the cancer cells was not due to mutations in Bcl-xL. This raised a possibility that Bcl-xL-interacting proteins involve in its subcellular localization changes.

### C-terminal Binding Protein 2 (CtBP2) is a novel Bcl-xL interacting protein

To identify Bcl-xL-interacting proteins that might promote its nuclear translocation, we employed an “ReCLIP” (Reversible Cross-Link Immuno-Precipitation) procedure using cell-permeable, thiol-cleavable crosslinkers to stabilize normally labile interactions *in situ* prior to isolation (Smith *et al*, 2011). We used human PNET BON1/TGL cells overexpressing HA-tagged wild-type (wt) Bcl-xL and the two HA-tagged Bcl-xL mutants defective in anti-apoptosis (mt1 and mt2) (Choi *et al*., 2016) for ReCLIP with anti-HA magnetic beads. The proteins in ReCLIP were subjected to mass spectrometry analysis to identify Bcl-xL interacting proteins. Among the specific proteins immunoprecipitated by anti-HA magnetic beads from the cells overexpressing HA-tagged wt Bcl-xL and Bcl-xL mutants, but not immunoprecipitated from the parental cells overexpressing the control vector, we focused on the proteins with both a nuclear localization signal (NLS) and a role in metastasis. The top candidate was CtBP2 (Supplementary Table 1) because of the following reasons. First, CtBP2 is a transcriptional regulator containing an NLS (Chinnadurai, 2003, 2009; Fang *et al*, 2006; Paliwal *et al*, 2012). It can act as a transcriptional repressor or activator. Second, CtBP2 is overexpressed in cancer, including prostate cancer, ovarian cancer, liver cancer, breast cancer, and esophageal squamous cell carcinoma (Cerami *et al*, 2012; Gao *et al*, 2013; Takayama *et al*, 2014; Zhang *et al*, 2015; Zheng *et al*, 2015). Third, CtBP2 has been shown to promote epithelial-mesenchymal transition (EMT) and migration of several types of cancer cells (Wang *et al*, 2013; Zhang *et al*., 2015; Zheng *et al*., 2015). Fourth, H3K4me3 is enriched at the CtBP2-binding sites in human embryonic stem cells (Lee *et al*, 2015) and Bcl-xL overexpression increases global H3K4me3 levels in cancer cells (Choi *et al*., 2016).

To verify the interaction between CtBP2 and HA-Bcl-xL identified by IP-mass spectrometry, we performed anti-HA ReCLIP using lysates from BON1/TGL cells overexpressing HA-tagged wt Bcl-xL or the vector, followed by Western blotting using antibodies against CtBP2 and HA. In consistent with the IP-mass spectrometry results, CtBP2 was detected in the anti-HA ReCLIP specifically from cells expressing HA-tagged wt Bcl-xL, but not from the control cells expressing the vector (Figure 2a).

**Figure 2.**
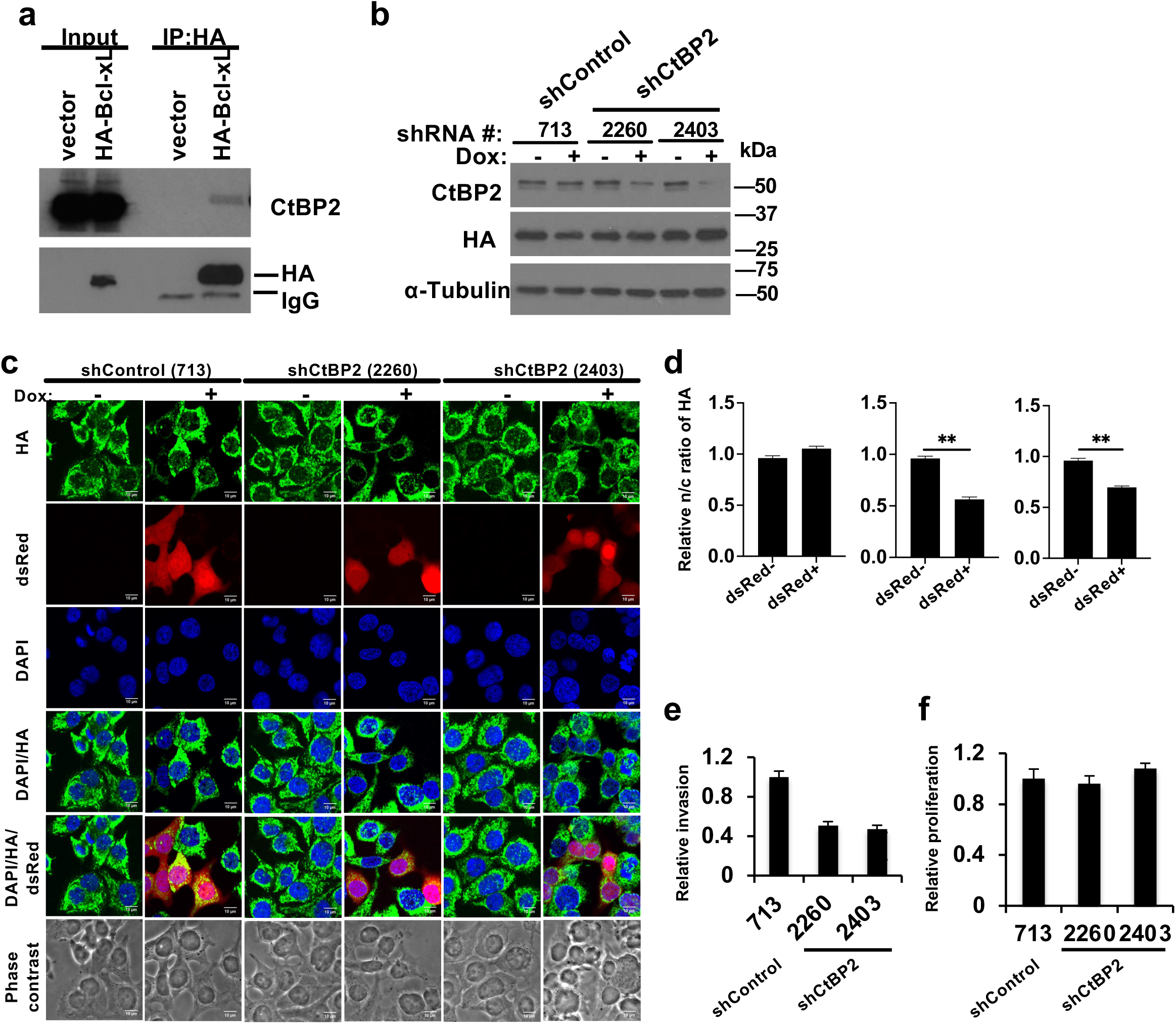
Identification of CtBP2 as a novel Bcl-xL interacting protein. (**a**) Whole cell lysates from BON1/TGL cells that stably express HA tagged Bcl-xL or pQCXIP (pQ) vector were used for ReCLIP using anti-HA magnetic beads and Western blotting using CtBP2 and HA anti-bodies. (**b**) BON1/TGL/pQ and BON1/TGL/HA-Bcl-xL cells expressing dox-inducible shCtBP2 or shRLuc were treated with 1 μg/ml doxycycline (dox) for 96 hours and harvested for Western blot analysis of CtBP2 and HA (for HA-Bcl-xL). α-tubulin was used a loading control. (**c**) Representative confocal images of BON1/TGL/HA-Bcl-xL/tet-O-dsRed-shRLuc (#713)-PGK-rtTA3, BON1/TGL/HA-Bcl-xL/tet-O-dsRed-shCtBP2 (#2260)-PGK-rtTA3, and BON1/TGL/HA-Bcl-xL/tet-O-dsRed-shCtBP2 (#2403)-PGK-rtTA3 cells that were treated or untreated with 1 μg/ml Dox for 96 hours, fixed, and stained with anti-HA antibody (green) and DAPI (blue). Scale bar, 10 μm. Original magnification, × 60. (**d**) The nuclear to cytosol (n/c) ratio of HA-Bcl-xL florescent signals were quantified in at least 40 individual cells per cell line and normalized to the dsRed negative. Results are reported as mean ± SEM. Data were analyzed by one-way ANOVA followed by Turkey HSD *post hoc* test. *: statistically significant difference at *p*<0.05. (**e** and **f**) Cells pre-treated with 0.5 μg/ml doxycycline for 96 hours were assayed for transwell invasion (e) and proliferation. (f) Values are mean ± SEM, N= 3. *: *p*<0.05, two tail t test.

### CtBP2 promotes the nuclear translocation of Bcl-xL

To determine the role of CtBP2 in mediating Bcl-xL’s promotion of migration, we employed a loss of function approach by knocking down *CTBP2* using doxycycline (dox)-inducible shRNA against *CTBP2* in BON1/TGL/HA-Bcl-xL cells. We constructed four dox-inducible shRNAs (#2260, #2403, #2255, and #2805) against different regions of human CtBP2 and a control shRNA against Renilla Luciferase (shRLuc #713) in a lentivirus vector, LT3RENIR (Fellmann *et al*., 2013) (Supplementary Table S2). This LT3RENIR vector expresses dsRed-coupled miR-E shRNA from an optimized Tet-responsive element promoter (T3G), which strongly reduces the leaky shRNA expression, and its miR-E backbone increases the mature shRNA levels and knockdown efficacy (Fellmann *et al*., 2013). It also harbors rtTA3 to enable single-vector (“all-in-one”) Tet-ON shRNAmir expression (Fellmann *et al*., 2013). We used the “tet-O-ds<u>R</u>ed-shRNA-PGK-rtTA3 (TR-shRNA)” viruses to infect BON1/TGL/HA-Bcl-xL cells (BON1/HA-Bcl-xL). Infected cells were selected with neomycin to obtain stable clones. We investigated knockdown efficiency of CtBP2 by these different shRNAs in the stable cell lines following 96 hours of doxycycline treatment and Western blotting. Two (#2260 and #2403) shRNAs against CtBP2 successfully reduced the levels of CtBP2 proteins (Figure 2b).

Co-expression of dsRed and shRNAs by dox induction allows dsRed reporter-based estimation of shRNA expression by a fluorescent microscope. We observed that ∼ 50% of cells showed positive dsRed signal after 1 μg/ml doxycycline treatment for 96 hours, indicating a mixed cellular response to doxycycline induction and thus shRNA expression and CtBP2 knockdown (Supplementary Figure S2). This gave us a unique opportunity to determine whether knockdown of CtBP2 affected Bcl-xL nuclear localization by comparing subcellular localization of HA-Bcl-xL in the dsRed-positive cells and the dsRed-negative cells in the same images. We have previously shown that the nuclear Bcl-xL, but not its cytoplasmic counterpart, accounts for its migration/invasion activity (Choi *et al*., 2016). Using anti-HA immunofluorescent staining, we observed similar nuclear/cytoplasmic ratio of HA-Bcl-xL in the dsRed-positive and dsRed-negative BON1/HA-Bcl-xL/TR-shRLuc #713 control cells (Figure 2c and 2d). In contrast, the nuclear/cytoplasmic ratio of HA-Bcl-xL was significantly reduced in the dsRed-positive shCtBP2 cells compared to the dsRed-negative cells in both BON1/HA-Bcl-xL/TR-shCtBP2 #2260 and #2403 cultures (Figure 2c and 2d), suggesting that knockdown of CtBP2 inhibits nuclear translocation of Bcl-xL.

### CtBP2 knockdown decreases the ability of Bcl-xL to promote invasion and metastasis

To examine whether the invasion function of Bcl-xL is mediated by CtBP2, we performed transwell invasion assays using dox-treated BON1/HA-Bcl-xL/TR-shCtBP2 #2260 and #2403 cells and BON1/HA-Bcl-xL/TR-shRLuc #713 control cells. We found that invasion activity was significantly reduced in dox-treated shCtBP2 (#2260 and #2403) cells compared with dox-treated shRLuc (#713) control cells (Figure 2e). On the other hand, there was no significant difference in cell proliferation among all three cells lines treated with doxycycline (Figure 2f). The data suggested that CtBP2 knockdown reduced Bcl-xL-dependent invasion of BON1/TGL/HA-Bcl-xL cells.

As approximately 50% of dox-treated cells showed positive dsRed signals, a reporter for shRNA induction by doxycycline (Supplementary Figure S2), we enriched shRNA-expressing cells by fluorescence-activated cell sorting (FACS) for dsRed-positive cells. dsRed was observed in >90% sorted cells by fluorescent microscopy (Figure 3a). It is of note that dsRed-positive cells gradually lost dsRed expression after long-term doxycycline treatment (data not shown). Therefore, we maintained and cryopreserved the sorted dsRed-positive cell lines without doxycycline and only started doxycycline treatment 96 hours before analyzing the effect of shRNAs. We detected significant dox-induced CtBP2 knockdown in the sorted, dsRed-positive BON1/HA-Bcl-xL/TR-shCtBP2 #2260 and BON1/HA-Bcl-xL/TR-shCtBP2 #2403 cells (Figure 3b).

**Figure 3.**
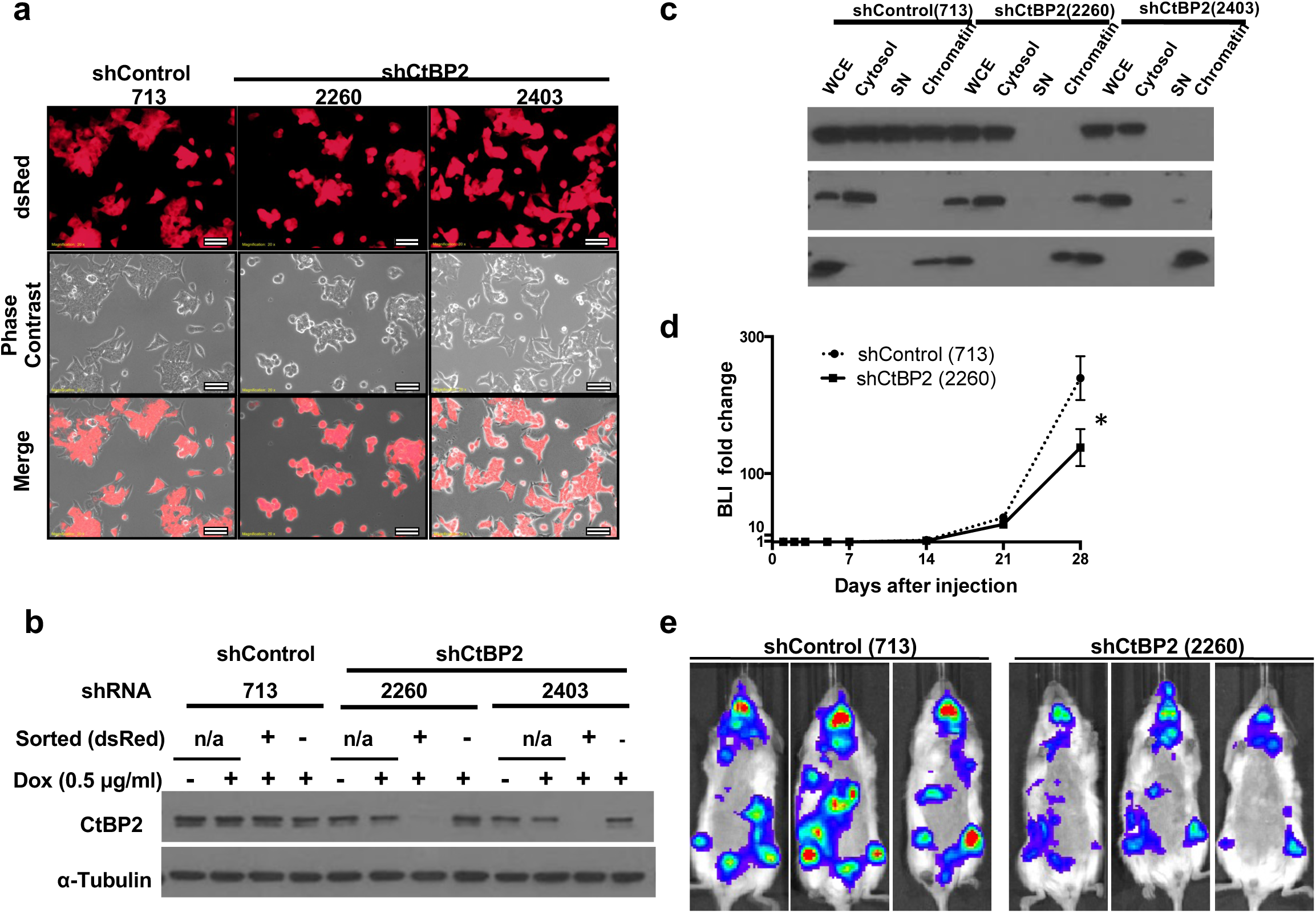
Knockdown of CtBP2 reduces metastasis promoted by Bcl-xL *in vivo*. (**a**) Representative fluorescent images of sorted, dsRed positive BON1/TGL/HA-Bcl-xL/tet-O-dsRed-shRLuc (#713)-PGK-rtTA3, BON1/TGL/HA-Bcl-xL/tet-O-dsRed-shCtBP2 (#2260)-PGK-rtTA3, and BON1/TGL/HA-Bcl-xL/tet-O-dsRed-shCtBP2 (#2403)-PGK-rtTA3 cells pre-treated with 0.5 μg/ml doxycycline for 96 hours. Scale bar, 50 μm. Original magnification, × 20. (**b**) Western blot analysis of CtBP2 protein levels. α-tubulin was used as loading control. (**c**) Cells were subjected to the biochemical fractionation. Mek1/2 (cytosolic protein) and histone H3 (nuclear protein) were used as controls. (**d**) Dox-pretreated BON1/TGL/HA-Bcl-xL/tet-O-dsRed-shRLuc (#713)-PGK-rtTA3 and BON1/TGL/HA-Bcl-xL/tet-O-dsRed-shCtBP2 (#2260)-PGK-rtTA3 cells were injected into NSG mice that started doxycycline diet one day before tumor injection. Fold change of bioluminescent signals over the course of the 28 days was shown. N= 5 for each group. (**e**) Representative bioluminescent images of mice at day 28 were shown from each group.

We next performed biochemical fractionation analysis using these FACS sorted, dsRed-positive cells and found that both soluble nuclear HA-Bcl-xL and chromatin-bound HA-Bcl-xL were greatly reduced in the dox-induced CtBP2 knockdown cells (#2260 and #2403) compared to the dox-induced shRLuc control cells (#713) (Figure 3c). Mek1/2 (cytoplasmic proteins) and histone H3 (a nuclear protein) were used in the Western blotting as the reference for each fractionation respectively (Figure 3c). Consistent with the immunofluorescence results, the parallel biochemical subcellular fractionation assay further validated that CtBP2 mediated the nuclear translocation of Bcl-xL.

As our data from clinical specimens and invasion assays suggested that CtBP2-driven nuclear-localization of Bcl-xL may promote metastasis, we performed experimental metastasis assays in mice using FACS sorted cells. We intracardiacally injected 1 x 10^6^ dox-treated BON1/HA-Bcl-xL/TR-shRLuc #713 (control cells) and BON1/HA-Bcl-xL/TR-shCtBP2 #2260 cells into NOD/scid-lL2Rgc knockout (NSG) immunodeficient mice. Mice were started on a doxycycline diet one day before tumor injection and kept on doxycycline diet until the end point. The luciferase expression in these cells provided a mean for monitoring the localization and growth of tumor cells through *in vivo* bioluminescent imaging. Indeed, weekly bioluminescent imaging showed that bioluminescence signals started to increase after 14 days, and metastatic cells were detected at multiple sites on day 28 (Figure 3d and 3e). The increase in bioluminescent signals over time from the CtBP2 knockdown group was significantly less than that from the control shRLuc (#713) group (Figure 3d, GEE analysis, *p* = 0.0366), suggesting that knockdown of CtBP2 reduced metastatic progression of Bcl-xL-overexpressing BON1 cells in mice.

### CtBP2 knockout reduces Bcl-xL transcripts and the nuclear pool of Bcl-xL

In addition to shRNA knockdown of CtBP2 in BON1 PNET cells, we used CRISPR-Cas9 to knock out (KO) CtBP2 in the MDA-MB-231 breast cancer cell line to further test the regulation of nuclear translocation of Bcl-xL by CtBP2. We found that CtBP1, a homolog of CtBP2, did not increase to compensate the loss of CtBP2, but CtBP2 KO reduced endogenous Bcl-xL protein levels (Figure 4a). Because CtBP2 is a transcription cofactor, we investigated the possibility that CtBP2 regulated Bcl-xL transcription. Our RT-PCR confirmed that CtBP2 mRNA was not detectable in the KO cells (Figure 4b) and Bcl-xL mRNA was significantly reduced in the CtBP2 KO cells compared to the control cells (Figure 4c). Furthermore, we searched the public research project website, The Encyclopedia of DNA Elements (ENCODE) (Consortium, 2004, 2011), and found that CtBP2 bound near the transcription start site of Bcl-xL in the dataset of CtBP2 ChIP-Seq from human H1-hESC cell line (Supplemental Figure S2). Taken together, CtBP2 also functions as a transcription activator of Bcl-xL.

**Figure 4.**
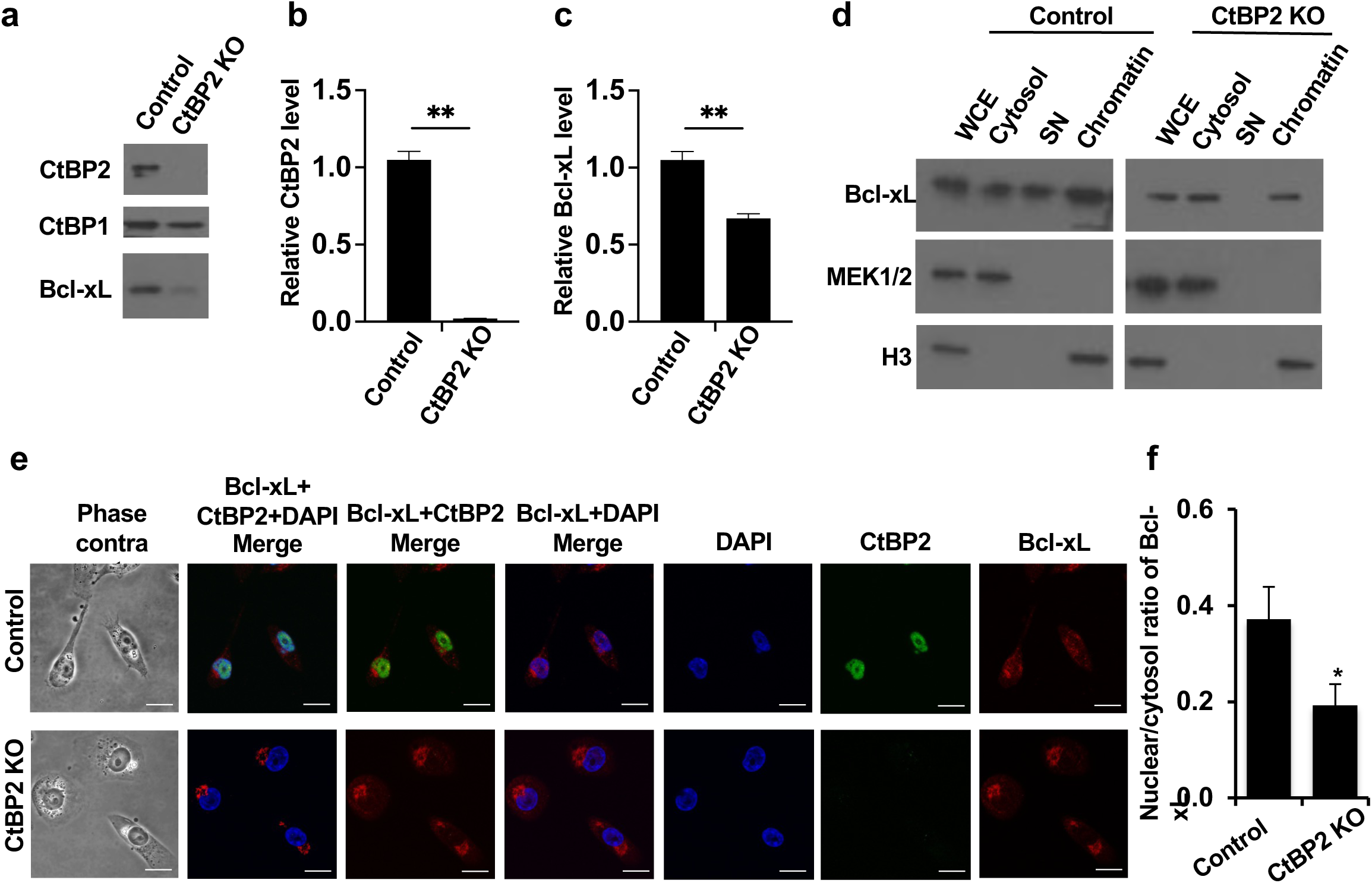
Knockout of CtBP2 reduces Bcl-xL transcripts and the nuclear to cytosol ratio of Bcl-xL. (**a**) MDA-MB-231 control cells and MDA-MB-231 with CtBP2 knockout (KO) cells were harvested for Western blot analysis of CtBP2, CtBP1, and Bcl-xL protein levels. (**b** and **c**) RT-qPCR analysis of the CtBP2 (**b**) and Bcl-xL (**c**) expression levels. (**d**) Cells were subjected to the biochemical fractionation. Mek1/2 (cytosolic protein) and histone H3 (nuclear protein) were used as controls for the biochemical fractionation. (**e**) Cells were fixed and stained with anti-CtBP2 antibody (green), anti-Bcl-xL antibody (red), and DAPI (blue). Representative confocal images were shown. Scale bar, 20 μm. Original magnification, × 60. (**f**) The nuclear to cytosol ratio of Bcl-xL florescent signals were quantified in at least 40 individual cells. Data were analyzed by one-way ANOVA followed by Turkey HSD *post hoc* test and shown as mean ± SEM. *: *p*<0.05.

To determine whether CtBP2 KO also reduced nuclear subcellular localization of endogenous Bcl-xL, we performed biochemical fractionation and immunocytochemical staining in CtBP2 KO cells. The biochemical fractionation analysis showed that the nuclear pool of endogenous Bcl-xL was greatly reduced in the CtBP2 KO cells (Figure 4d). Immunofluorescent staining using antibodies against endogenous Bcl-xL and CtBP2 showed that the Bcl-xL signals were weaker in CtBP2 KO cells than in the control cells and the nuclear/cytoplasmic ratio of Bcl-xL in CtBP2 KO cells was significantly reduced (Figure 4e and 4f). Taken together, CtBP2 plays a dual role of regulating Bcl-xL transcription and translocating Bcl-xL proteins into the nucleus, both of which contribute to promote Bcl-xL’s pro-metastatic activity.

### The residues 82 to 104 of CtBP2 and the N-terminus of Bcl-xL are required for their interaction

To characterize the Bcl-xL/CtBP2 interaction domains, we used deletion constructs of CtBP2 to define the Bcl-xL interaction domain of CtBP2 (Figure 5a). DNA constructs of 3 different V5-tagged CtBP2 constructs (Full length (445 aa), residues 1 to 321, and residues 322 to 445) (Paliwal *et al*., 2006) and HA-tagged full-length Bcl-xL were transiently co-expressed in U2OS cells. After 48 hours, cell lysates were prepared for anti-HA ReCLIP. We found that full length CtBP2 and N-terminal CtBP2 (residues 1 to 321), but not C-terminal CtBP2 (residues 322 to 445) bound to HA-Bcl-xL (Figure 5b).

**Figure 5.**
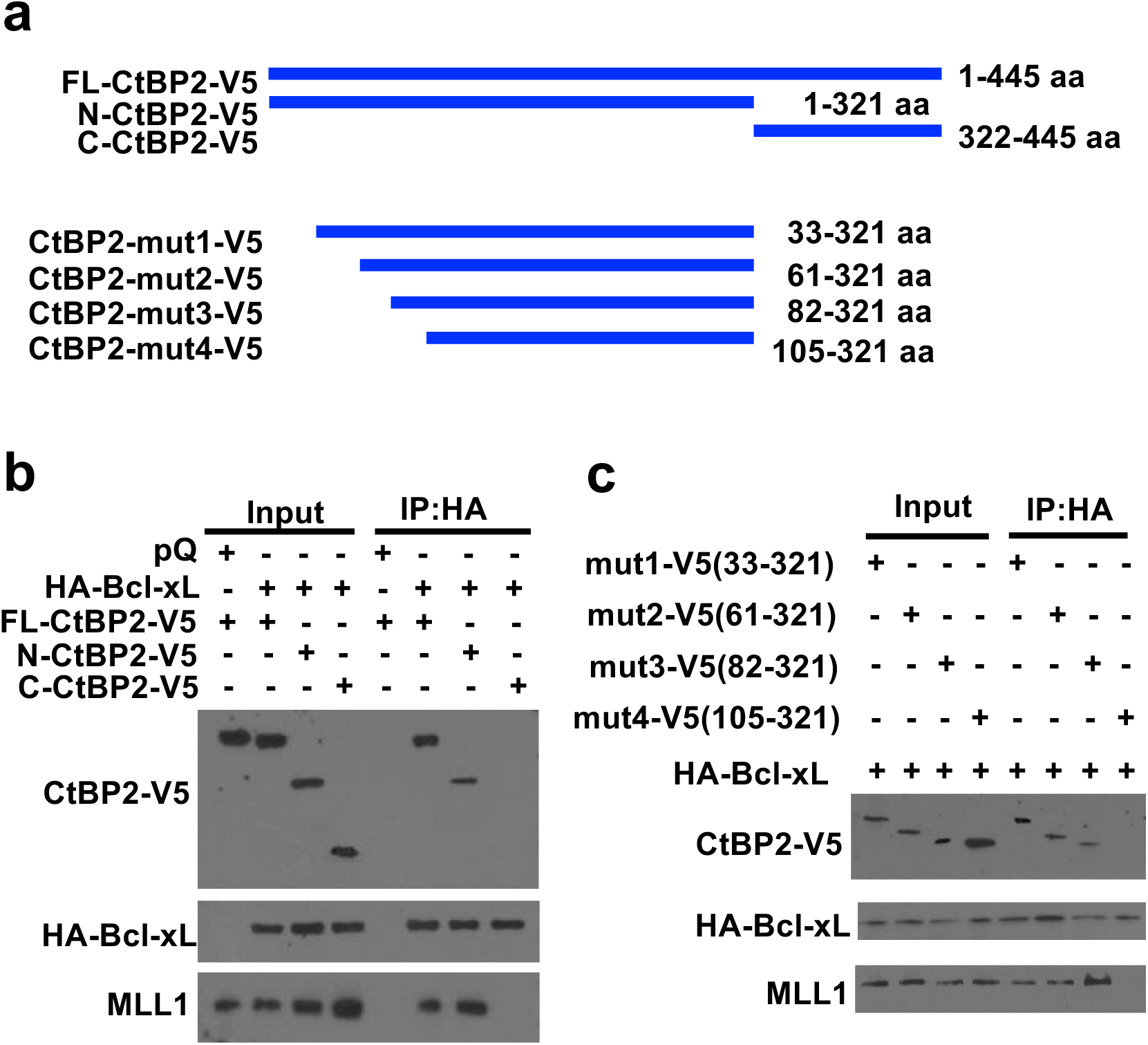
Mapping the Bcl-xL/CtBP2 interaction domain on CtBP2. (**a**) Schematic diagrams of full-length CtBP2 and its truncation constructs. (**b** and **c**) V5 tagged full length (FL) CtBP2 or CtBP2 truncation constructs were co-transfected with HA-tagged Bcl-xL to U2OS cells. 500 μg of clarified U2OS lysates were used for ReCLIP with anti-HA agarose followed by Western blotting for V5, HA, and MLL1.

To further narrow down the Bcl-xL-binding domain in the N-terminal region of CtBP2, we generated a series of CtBP2 N-terminal deletion constructs with a V5 tag (Figure 5a, mut1: residues 33 to 321, mut2: residues 61 to 321, mut3: residues 82 to 321, and mut4: residues 105 to 321) to teste for their capability to bind Bcl-xL. DNA constructs of HA-tagged Bcl-xL and these V5-tagged CtBP2 deletion constructs were transiently co-expressed in U2OS cells. After 48 hours, cell lysates were prepared for anti-HA ReCLIP. CtBP2 deletion constructs #1-3, but not #4, were detected in the HA-Bcl-xL IP (Figure 5c), suggesting that residues 82 to 104 of CtBP2 were required for its interaction with Bcl-xL.

To determine the CtBP2-binding domain in Bcl-xL, we performed the reciprocal experiments. We initially generated a series of N-terminal deletion Bcl-xL mutants with HA tag, but they did not express well (data not shown). Because we found that Bcl-B, another Bcl-2 anti-apoptotic family member (Beverly & Varmus, 2009), did not interact with V5-tagged CtBP2 in anti-V5 ReCLIP (Figure 6a and 7b, construct #1), we utilized a set of chimeric Bcl-xL/Bcl-B proteins (Saurabh *et al*., 2014) to narrow down the CtBP2 binding domain in Bcl-xL. Bcl-xL (#1), Bcl-B (#2), and six chimeric Bcl-xL/Bcl-B constructs (#3-8) were transfected into 293T cells, and all could be transiently expressed to similar levels (Figure 6b). Anti-V5 (for CtBP2) ReCLIP revealed that WT Bcl-xL, construct #4 that contains Bcl-xL’s BH4 domain, BH3 domain, and two N-terminal loop domain and construct #5 that contains Bcl-xL’s BH4 domain and the first N-terminal loop domain were detected in the anti-V5 IP, but the other chimeric proteins were not detected in the anti-V5 IP (Figure 6b). Therefore, the BH4 domain and the first N-terminal loop domain (residues 1 to 85) of Bcl-xL were needed for its interaction with CtBP2.

**Figure 6.**
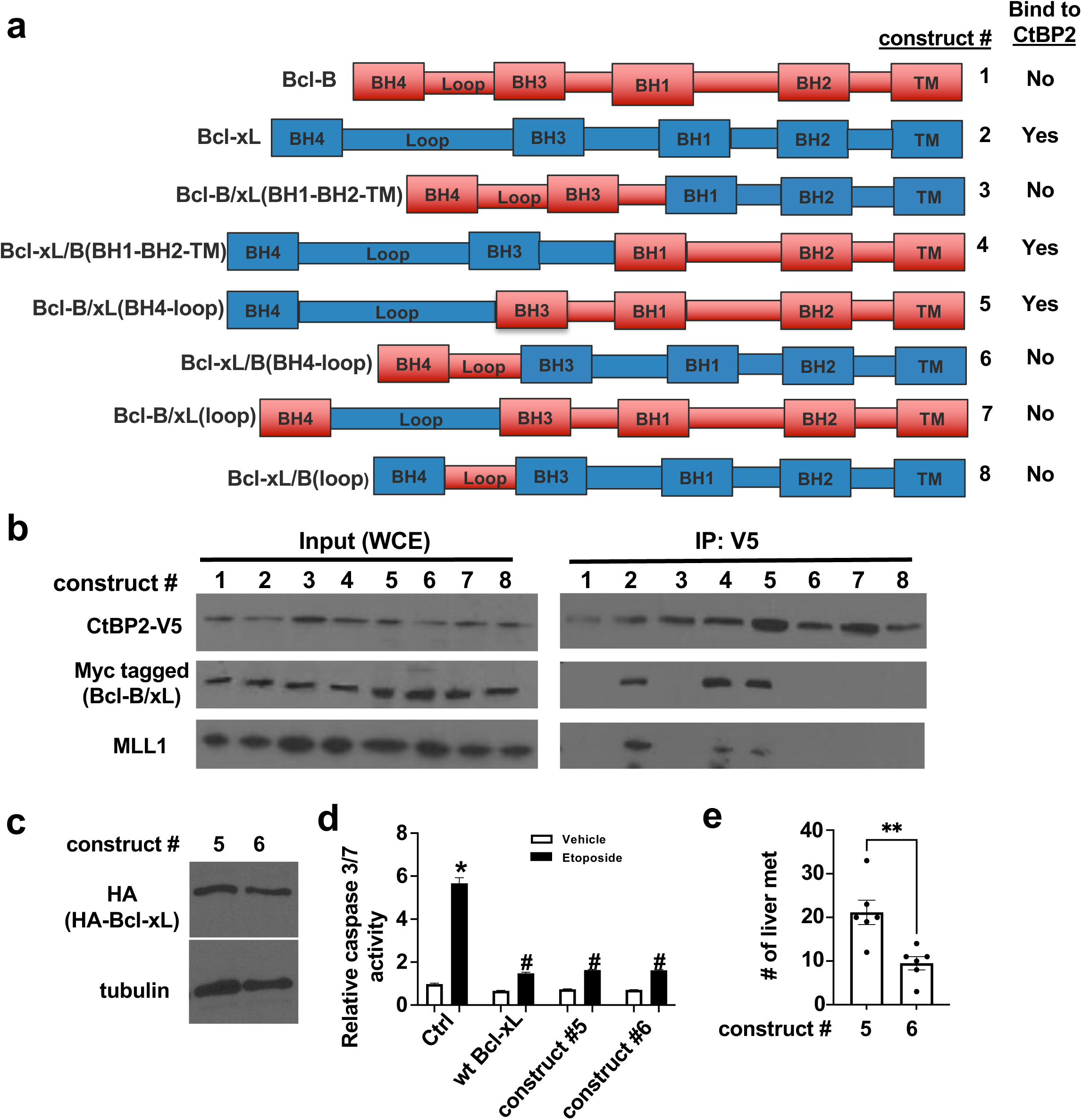
Mapping the Bcl-xL/CtBP2 interaction domain on Bcl-xL. (**a**) Schematic diagrams of chimeric Myc-tagged Bcl-xL/Bcl-B. (**b**) V5 tagged full length CtBP2 or Myc-tagged Bcl-xL/Bcl-B were co-transfected to 293T cells. The cell lysates were used for anti-Myc IP followed by Western blotting for V5, Myc, and MLL1. (**c**) Construct # 5 and 6 subcloned into HA-tagged RCAS vectors and were expressed in N134 cells. The expression of HA chimeric proteins was verified by Western blot. (**d**) Indicated cell lines were treated 10 mM etoposide or vehicle. After 24h, apoptosis was measured by Caspase-Glo 3/7 Assay. Results were presented as mean ± the standard error of the mean (SEM). *: *p*<0.05 compared with vehicle-treated control cells and #: *p*<0.05 compared with etoposide - treated control cells, student’s t-test. (**e**) N134-HA-construct #5 and N134-HA-construct #6 cells were injected into tail vein of NSG mice. Five weeks later, liver metastases (> 1 mm in diameter on Hematoxylin and eosin (H&E)-stained slides) were quantified.

**Figure 7.**
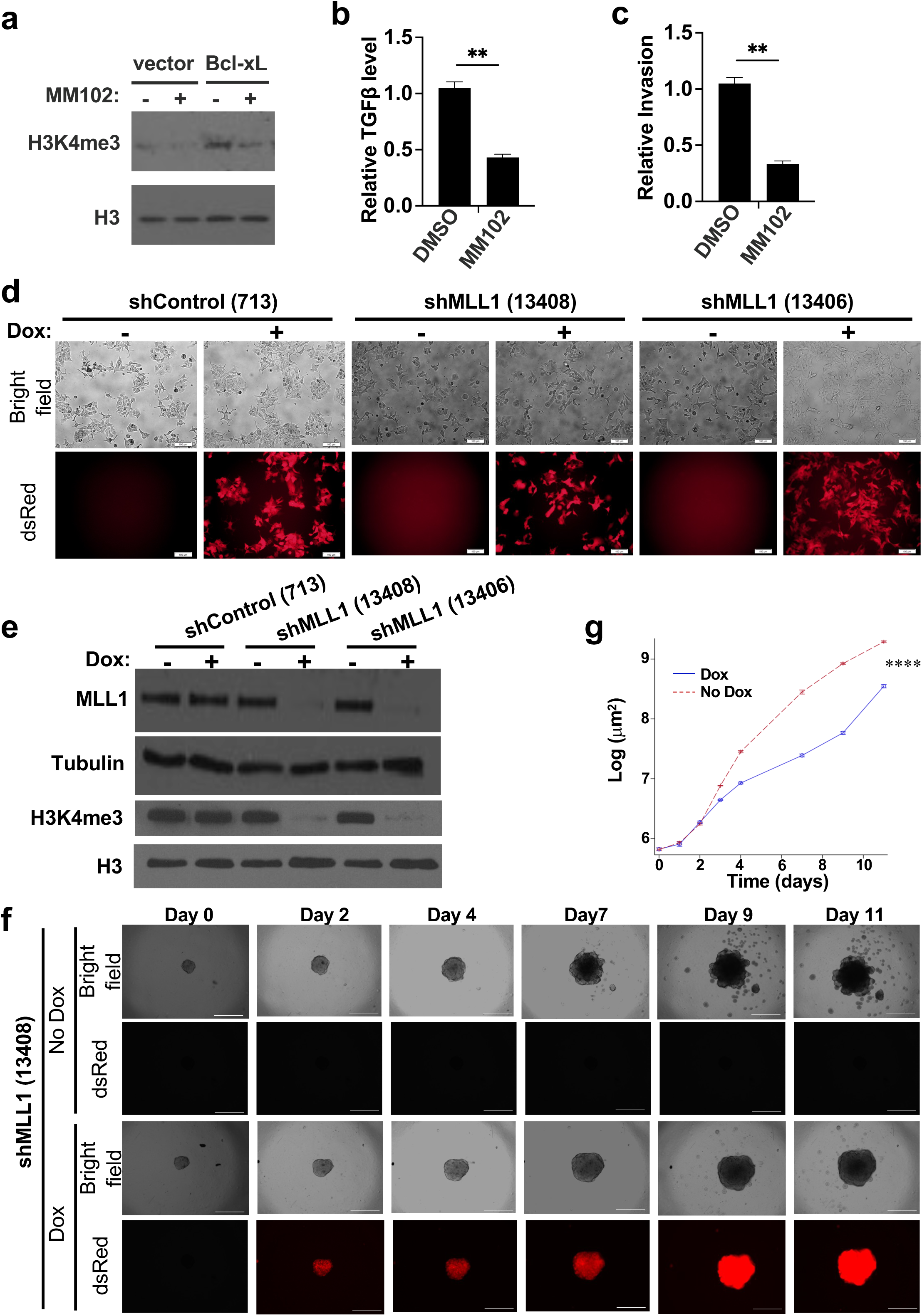
MLL1 interacts with the Bcl-xL/CtBP2 complex and knockdown of MLL1 reduces tumor spheroid invasion. (**a**) BON1/TGL cells overexpressing pQ vector or HA-Bcl-xL were treated with DMSO or 40 μM MM102 (MLL1 inhibitor) for 96 hours with medium changing for every 48 hours and harvested for Western blotting for H3K4me3 and total histone H3. (**b**) TGFβ mRNA levels in the BON1/TGL cells overexpressing Bcl-xL were analyzed by RT-qPCR. (**c**) Transwell invasion assay of the BON1/TGL cells overexpressing Bcl-xL treated with DMSO or MM102 was performed. Data are expressed as the normalized number of cells invaded to the bottom surface of the transwell inserts in eight fields under × 20 magnification after 24 hours relative to that of cells with vector alone. Values are means ± SEM. N= 3. \**p*<0.05, two tail t test. (**d**) Representative fluorescent images of BON1/TGL/HA-Bcl-xL/tet-O-dsRed-shRLuc (#713)-PGK-rtTA3, BON1/TGL/HA-Bcl-xL/tet-O-dsRed-shMLL1 (#13408)-PGK-rtTA3, and BON1/TGL/HA-Bcl-xL/tet-O-dsRed-shMLL1 (#13406)-PGK-rtTA3 cells treated with 1 μg/ml Dox for 96 hours were shown. Scale bar, 50 μm. Original magnification, × 20. (**e**) Cells were harvested for Western blotting of MLL1, H3K4me3 levels, α-tubulin and H3. (**f**) Representative images of spheroids of BON1/TGL/HA-Bcl-xL/tet-O-dsRed-shMLL1 (#13408)-PGK-rtTA3 cells. Original magnification ×4, scale bar 100 μm. (**g**) Change of whole tumor spheroid area over time was quantified (N= 6 for each condition). ****: *p*<0.0001, GEE method.

**Figure 8.**
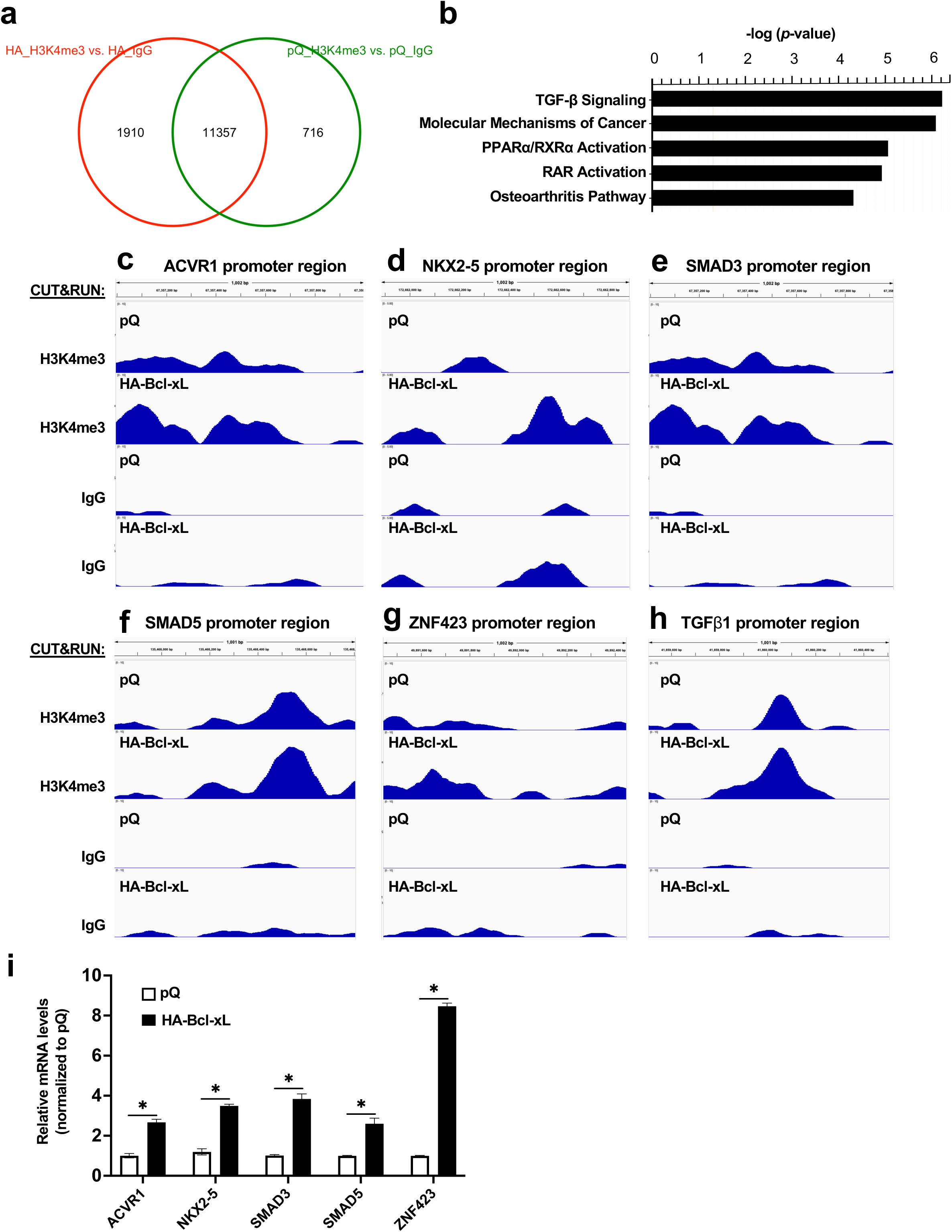
Bcl-xL positively regulates TGFβ signaling through H3K4me3 epigenetic modifications. (**a**) CUT&RUN-Seq analysis using anti-H3K4me3 antibody or IgG control in BON1/TGL/pQ (pQ) and BON1/TGL/HA-Bcl-xL (HA) cells was performed. Venn diagram of the number of H3K4me3 binding sites was shown. (**b**) IPA analysis of the gene promoters with unique H3K4me3 binding sites in BON1/TGL/HA-Bcl-xL cells. (**c-h**) Track views of representative Integrative Genomics Viewer (IGV) for peaks near the transcription start sites of ACVR1, NKX2-5, SMAD3, SMAD5, ZNF42, and TGFβ1 in CUT&RUN assays were shown. (**i**) RT-qPCR of ACVR1, NKX2-5, SMAD3, SMAD5, and ZNF42 from BON1/TGL/pQ and BON1/TGL/HA-Bcl-xL cells.

### The interaction between Bcl-xL and CtBP2 is important for the metastatic function of Bcl-xL

To investigate the relationship between Bcl-xL’s binding ability to CtBP2 and the metastatic function, we performed tail vein experimental metastasis assays. We sub-cloned two chimeric constructs, #5 (which binds to CtBP2) and #6 (which does not bind to CtBP2) into an avian retroviral vector, RCASBP, with an HA tag. We infected the N134 cell line (Du *et al*., 2007), which is derived from an PNET in a *RIP-Tag; RIP-tva* mouse, with RCASBP–HA-Bcl-B/xL(BH4-loop) (construct #5) or RCASBP–HA-Bcl-xL/B(BH4-loop) (construct #6). The expression levels of both HA-tagged chimeric proteins in N134 cells were similar by Western blot analysis (Figure 6c). Both chimeric constructs #5 and #6 protected cells from apoptosis as effectively as the wild-type Bcl-xL (Fig. 7d). We injected 1.5 x 10^6^ N134 cells expressing chimeric constructs #5 and #6 into the tail vein of NSG mice. After five weeks, organs of the recipient mice were harvested to survey for metastases. We found significantly more liver macrometastases in recipients of construct #5 that binds to CtBP2 than construct #6 that does not bind to CtBP2 (Figure 6e and Supplemental Figure S3), suggesting that the interaction between Bcl-xL and CtBP2 is important for the metastatic function of Bcl-xL.

### Mixed lineage leukemia protein-1 (MLL1) interacts with the CtBP2 and Bcl-xL complex

We previously reported that Bcl-xL increases H3K4me3 and the levels of TGFβ1 through H3K4me3 epigenetic modification at the TGFβ1 promoter (Choi *et al*., 2016). However, the mechanism by which Bcl-xL increases H3K4me3 was unknown. Because H3K4 methyltransferase, MLL1 (also known as KMT2A), has a putative CtBP2 binding motif (PXDLS), we hypothesized that the Bcl-xL/CtBP2 complex recruits MLL1 to epigenetically regulate transcription of key genes in metastasis. To test this, we investigated whether MLL1 is present in the Bcl-xL/CtBP2 protein complex. We detected the presence of MLL1 in the anti-HA (for Bcl-xL) or anti-V5 (for CtBP2) ReCLIP only when transiently co-expressing the constructs of Bcl-xL and CtBP2 that were able to interact with each other in U2OS cells (Figure 5b, 5c, and 6b). We were not able to detect MLL1 in the ReCLIP when the constructs of Bcl-xL and CtBP2 did not interact with each other in U2OS cells (Figure 5b, 5c, and 6b). Furthermore, we performed anti-HA ReCLIP using lysates from BON1/TGL cells overexpressing HA-Bcl-xL or vector, and we detected MLL1 and CtBP2 in the HA ReCLIP from BON1/TGL cells overexpressing HA-Bcl-xL (Supplemental Figure S4). Altogether, the data suggest that the interaction between CtBP2 and Bcl-xL is required for the binding of MLL1 into this protein complex.

To investigate whether the histone methyltransferase activity of MLL1 is required for Bcl-xL to raise H3K4me3 levels, increase TGFβ1 transcription, and promote invasion, we treated cells with MM-102, a high-affinity, small-molecule peptidomimetic MLL1 inhibitor (Karatas *et al*, 2013). As previously reported (Choi *et al*., 2016), overexpression of Bcl-xL increased total H3K4me3 levels (Figure 7a, lane 3). We found that treatment of MM-102 significantly reduced total H3K4me3 levels (Figure 7a), decreased TGFβ1 transcripts (Figure 7b), and suppressed invasion induced by Bcl-xL in transwell invasion assays (Figure 7c).

In addition to pharmacological inhibition of MLL1 activity, we generated dox-inducible shRNA to silence MLL1. We designed six different shRNAs against human MLL1 with the dsRed reporter (Supplementary Table S2) and generated stable BON1/TGL/HA-BcL-xL/tet-O-dsRed-shMLL1-PGK-rtTA3 (BON1/HA-Bcl-xL/TR-shMLL1) cell lines. After 96-hour doxycycline treatment, almost all cells became dsRed-positive (Figure 7d). We examined the knockdown efficacy of MLL1 by Western blot analysis. We found that two shMLL1 constructs (#13408 and #13406) led to the most effective MLL1 knockdown and to a drastic reduction of H3K4me3 levels (Figure 7e). To determine whether MLL1 mediates the metastatic function of Bcl-xL *in vitro*, we performed a 3D tumor spheroid invasion assay. We seeded BON1/HA-Bcl-xL/TR-shMLL1 #13408 cells into an ultra-low attachment plate with Matrigel. 96 hours later (labeled as Day 0 in Figure 7f and 7g), we began treatment with 0.5 μg/ml doxycycline. We found that knockdown of MLL1 in the dox group suppressed tumor spheroid invasion of the Bcl-xL-overexpressing cells (Figure 7f). GEE analysis of whole invasive area changes over time showed that the dox group (shMLL1) was significantly less than the no dox group (*p*<0.0001) (Figure 7g). On the other hand, knockdown of the control RLuc did not reduce tumor spheroid invasion (Supplemental Figure S5).

### TGFβ signaling is the top pathway promoted by Bcl-xL-induced H3K4me3 histone modifications

To detect the sites of H3K4me3 histone modification by Bcl-xL, we performed Cleavage Under Targets and Release Using Nuclease (CUT&RUN) assay (Skene & Henikoff, 2017) on the BON1/TGL cells overexpressing HA-Bcl-xL (BON1/HA-Bcl-xL) and the control cells (BON1/pQ) with anti-H3K4me3 antibodies. Next-generation sequencing (NGS) revealed 1,190 unique H3K4me3 histone modification regions in the BON1/HA-Bcl-xL cells but not in the control BON1/pQ cells (Fig. 9a). Using Ingenuity Pathway Analysis (IPA) for these 1,190 unique H3K4me3 histone modification regions, we found that the top canonical pathway was TGFβ signaling (Fig. 9b). Fig. 9c-9g showed the H3K4me3 and IgG peaks around the transcription start sites of ACVR1, NKX2-5, SMAD3, SMAD5, and ZNF42 that belong to the TGFβ signaling pathway. The CUT&RUN-Seq also revealed differential enrichment of H3K4me3 histone modifications in the TGFβ1 promoter region in the BON1/HA-Bcl-xL cells compared to the control pQ cells (Fig. 9h). RT-qPCR showed that mRNA levels of ACVR1, NKX2-5, SMAD3, SMAD5, and ZNF42 were upregulated in HA-Bcl-xL overexpressing cells compared to control pQ cells (Fig. 9i), suggesting a correlation between the transcription and the increased H3K4me3 marks identified in the CUT&RUN-Seq assay.

## Discussion

Metastasis, the dissemination of cancer cells from primary sites to distant organs, accounts for the majority of cancer-associated mortality. Efforts on unraveling the molecular basis of tumorigenesis and metastasis have significantly advanced our understanding of metastasis. During cancer progression, the ability to override apoptosis is a prominent mechanism that cancer cells usually acquire (Hanahan & Weinberg, 2011). The Bcl-2 protein family plays a central role in regulating the intrinsic apoptosis pathway at the outer mitochondria membrane. At least twelve structure-related members of the Bcl-2 superfamily have been identified which could be categorized to three groups: the pro-apoptotic Bax and Bak, anti-apoptotic members like Bcl-2 and Bcl-xL, and BH-3 only proteins that cooperate with Bax and Bak to promote apoptosis (Youle & Strasser, 2008). Attempts at targeting anti-apoptotic Bcl-xL members in cancers have been primarily focused on inhibiting its anti-apoptotic activity, typically by disrupting its association with pro-apoptotic members and BH3-only proteins to release the sequester of these death-inducing factors and hence to amplify apoptotic signals. These strategies, however, showed limited success in clinical trials (Gandhi *et al*, 2011; Rudin *et al*, 2012), suggesting unrecognized alternative mechanisms for Bcl-xL to promote cancer progression.

This study, together with our previous findings (Choi *et al*., 2016), identifies a novel mechanism that CtBP2 transport overexpressed Bcl-xL into the nucleus and nuclear Bcl-xL drives metastasis of cancer cells independent of its anti-apoptotic activity. Nuclear import of macromolecules is a selective process. Proteins typically rely on the presence of NLS motifs to be recognized by specific transport receptors of the importin/karyopherin family, which mediate the transport of macromolecular cargos across the nuclear pore complex into the nucleus (Freitas & Cunha, 2009). Proteins without NLS sequences, like Bcl-xL, can be recruited to the nucleus through a “piggy-back” mechanism by associating with NLS-containing binding partners. Using a special “ReCLIP” technique to stabilize labile protein-protein interactions *in situ*, we discovered that CtBP2, a transcriptional regulator, translocates Bcl-xL into the nucleus. CtBP2 contains an KRQR NLS sequence at the N-terminal domain, which is crucial for its accumulation and gene repression functions in the nucleus (Verger *et al*, 2006; Zhao *et al*, 2006). In addition, CtBP2 shuttles between the nucleus and cytoplasm, and thus has been implicated in intracellular trafficking (Verger *et al*., 2006). We demonstrated that knockdown of CtBP2 significantly reduced the nuclear/cytoplasm partitioning of Bcl-xL and invasion and metastasis of the cancer cells induced by Bcl-xL overexpression *in vivo*.

The BH1, BH2, and BH3 domains of Bcl-xL create a hydrophobic groove to bind BH3-only pro-apoptotic proteins, inhibit their activity and lead to a pro-survival phenotype (Lee & Fairlie, 2019). The first loop domain of Bcl-xL is unstructured based on NMR or X-ray crystallographic analyses, and it does not contribute to anti-apoptotic activity of Bcl-xL (Muchmore *et al*, 1996). Here we showed that the BH4 domain and the first N-terminal loop domain (residues 1 to 85) of Bcl-xL are important for its interaction with CtBP2. The result of our domain mapping is in consistent with the finding that ABT-737, a BH-3 mimetic, did not affect the ability of Bcl-xL to promote cell migration (Choi *et al*., 2016). Because we found that another anti-apoptotic BCL family member, Bcl-B, did not bind to CtBP2, the interaction between Bcl-xL and CtBP2 is not shared among all anti-apoptotic BCL family members. On the other hand, we identified that residues 82 to 104 of CtBP2, which lie within a domain that recognizes PxDLS motif in many transcription factors that bind to CtBP2 (Dcona *et al*, 2017), are required for CtBP2 binding to Bcl-xL. Although CtBP1 interacts with the epigenetic regulator MLL1 (Xia *et al*, 2003), it was previously unknown whether CtBP2 interacts with MLL1. Here we showed that MLL1 associated with the Bcl-xL/CtBP2 complex. Despite containing a PxDLS motif, MLL1 only interacts with CtBP2 when Bcl-xL is present. It is likely that Bcl-xL causes exposure of the PxDLS motif in MLL1. We also postulate that MLL1 links to the Bcl-xL/CtBP2 complex to DNA/H3K4me3, and CtBP2 and Bcl-xL change MLL1’s H3K4me3 site selection as MLL1 fusion proteins are known to change transcriptional targets (Krivtsov *et al*, 2017; Xu *et al*, 2016). Further study will be required to identify the exact components of the Bcl-xL/CtBP2/MLL1 protein complex.

Enhanced TGFβ signaling promotes cancer metastasis (Derynck *et al*, 2021). We previously demonstrated that overexpression of Bcl-xL induces EMT and increased TGFβ production in a Bax/Bak-independent manner (Choi *et al*., 2016; Du *et al*., 2007). We also showed that Bcl-xL increased H3K4me3 at the promoter region of TGFβ and TGFβ-neutralizing antibodies significantly reduced Bcl-xL-mediated invasion (Choi *et al*., 2016). This work identified a Bcl-xL/CtBP2 complex that interacts with MLL1 to increase H3K4me3 modification on TGFβ1 and other genes involved in TGFβ signaling. Using MM102, a high-affinity, small-molecule peptidomimetic inhibitor for MLL1’s enzymatic activity, we demonstrated that H3K4 HMT activity of MLL1 is important for Bcl-xL to promote global levels of H3K4me3, TGFβ1 transcripts, and invasion. CtBP2 not only controls Bcl-xL nuclear translocation, but also regulates the transcription of Bcl-xL. It is likely that Bcl-xL drives CtBP2 transcription program as part of the CtBP2 transcription supercomplex to promote metastasis. Our work suggests a novel therapeutic perspective by targeting the interacting interface between CtBP2 and Bcl-xL, and thereby reducing the accumulation of Bcl-xL in the nucleus and attenuating metastasis progression.

## Acknowledgments

We thank Dr. Levi J. Beverly for the Bcl-xL/Bcl-B chimeric constructs (Saurabh *et al*., 2014). We thank Cheryl Zhang, George Zhang, Megan Wong, Tuo Zhang, Jiayi Liu, Makheni Jean-Pierre, Annie Yang, Dr. Sandra J. Shin, Dr. Yan Feng, Danny Huang, Dr. Hediye Erdjument-Bromage at MSKCC Microchemistry & Proteomics Core, Dr. Ralph Garippa, Qing Xiang, Ayana Suber, Hsiu-Yu Liu at MSKCC Gene Editing & Screening Core, Dr. Ruifang Li, Zhifan Yang, Nikita Albanese, and Dr. Pierre-Jacques Hamard at MSK Center for Epigenetics Research, and Dr. Bing He, Leticia Dizon, Mai Ho, Taotao Zhang, Ruben Diaz, Vincent Sarno at the Center for Translational Pathology at the Department of Pathology and Laboratory Medicine, Weill Cornell Medicine. This work is partially supported by DOD W81XWH-16-1-0619 (to Y.-C.N.D.), W81XWH-16-1-0620 (to Y.L.), and NIH 1R01CA204916 (to Y.-C.N.D.). YCND is The Rasweiler Family Research Scholar in Cancer Research.

## Author contributions

T.Z., S.L, J.N, X.C., S.C. designed and conducted the experiments and analyzed the data. A.Y.T., Z.C., D.W. performed bioinformatics analysis and statistical analysis. B.M. assisted with design and provided analysis tools for CUT&RUN. P.D. and S.R.G. generated CtBP2 deletion constructs and CtBP2 knockout MDA-MB-231 cell line. Y.-T.C. contributed to pathological characterization and molecular pathology. X.J. and Y.L. provided data interpretation. J.Z.X. supervised NGS. Y.-C.N.D. provided study conceptualization and supervision and wrote the manuscript. All authors read the manuscript, agreed with the content, and were given the opportunity to provide input.

## Availability of supporting data

The CUT&RUN-Seq datasets of this study are available in Gene Expression Omnibus (accession #GSE221629, https://www.ncbi.nlm.nih.gov/geo/query/acc.cgi?acc= GSE221629), and the other data are available from the authors upon reasonable request.

